# Lineage Commitment of Dermal Fibroblast Progenitors is Mediated by Chromatin De-Repression

**DOI:** 10.1101/2023.03.07.531478

**Authors:** Quan M. Phan, Lucia Salz, Sam S. Kindl, Jayden S. Lopez, Sean M. Thompson, Jasson Makkar, Iwona M. Driskell, Ryan R. Driskell

**Affiliations:** School of Molecular Biosciences, Washington State University, Pullman, WA; Center for Reproductive Biology, Washington State University, Pullman, WA; North Rhine-Westphalia Technical University of Aachen, Aachen, Germany

## Abstract

Dermal Fibroblast Progenitors (DFPs) differentiate into distinct fibroblast lineages during skin development. However, the mechanisms that regulate lineage commitment of naive dermal progenitors to form niches around the hair follicle, dermis, and hypodermis, are unknown. In our study, we used multimodal single-cell approaches, epigenetic assays, and allografting techniques to define a DFP state and the mechanisms that govern its differentiation potential. Our results indicate that the overall chromatin profile of DFPs is repressed by H3K27me3 and has inaccessible chromatin at lineage specific genes. Surprisingly, the repressed chromatin profile of DFPs renders them unable to reform skin in allograft assays despite their multipotent potential. Distinct fibroblast lineages, such as the dermal papilla and adipocytes contained specific chromatin profiles that were de-repressed during late embryogenesis by the H3K27-me3 demethylase, Kdm6b/Jmjd3. Tissue-specific deletion of Kdm6b/Jmjd3 resulted in ablating the adipocyte compartment and inhibiting mature dermal papilla functions in single-cell-RNA-seq, ChIPseq, and allografting assays. Altogether our studies reveal a mechanistic multimodal understanding of how DFPs differentiate into distinct fibroblast lineages, and we provide a novel multiomics search-tool within skinregeneration.org.

## INTRODUCTION

Fibroblasts are mesenchymal cells that create and maintain the extracellular matrix of the connective tissues supporting the functions of various organs (Driskell and Watt, 2015; Plikus et al., 2021; Watt and Fujiwara, 2011). They are distinct across different organs but are also diverse within the same tissue (Buechler et al., 2021; Dhouailly and Sengel, 1975; Fernandes et al., 2004; Muhl et al., 2020). Mammalian skin serves as a critical model to study development, homeostasis, and the role of fibroblast heterogeneity in tissue-resident stem cell niche, tissue regeneration, fibrosis-associated diseases, and the effects of chronic aging (Chen et al., 2013; Cole et al., 2018; Griffin et al., 2020; Gruber et al., 2020; Gurtner et al., 2008; Singer and Clark, 1999). Recent studies have demonstrated the distinct roles of dermal fibroblast subpopulations during embryonic development and skin wound healing. These studies have revealed that manipulations of specific fibroblast subpopulations lead to hair follicles and adipocytes regeneration in different wounding scenarios (Abbasi et al., 2020; Correa-Gallegos et al., 2019; Guerrero-Juarez et al., 2019; Jiang et al., 2018; Mascharak et al., 2021; Phan et al., 2020; Plikus et al., 2017; Rinkevich et al., 2015; Shook et al., 2020; Sinha et al., 2022). However, the establishment of fibroblast heterogeneity from a common progenitor during embryonic development remains poorly characterized.

During early embryonic development, mesenchymal stem cells migrate from different embryonic origins to sub-epidermal regions of the skin, where the dorsal dermis arises from the somite, and the facial dermal fibroblasts derive from the neural crest (Atit et al., 2006; Dhouailly, 1973; Fernandes et al., 2004; Thulabandu et al., 2018; Tran et al., 2010; Wong et al., 2006). At roughly E13.5-14.5 in murine skin, reciprocal signaling between the overlying epidermis and the Dermal Fibroblast Progenitors (DFPs) leads to condensation of upper fibroblasts to form the placodes (Chen et al., 2012; Fu and Hsu, 2013; Kishimoto et al., 2000; Millar, 2002). Between E14.5 and E16.5, the DFPs then differentiate and commit to distinct lineages of upper papillary and lower reticular fibroblasts (Driskell et al., 2013). Upper papillary fibroblasts differentiate into the dermal papilla, neonatal papillary fibroblast, and the arrector pili muscles post-natal. Lower reticular fibroblasts differentiate into collagen-secreting reticular and the dermal white adipocyte (DWAT)(Festa et al., 2011; Wojciechowicz et al., 2013). Moreover, classic chimeric skin grafting studies between avian and murine skin have shown that DFPs need to differentiate before responding to epidermal signals, and that fibroblast identity is also region-specific (Dhouailly, 1973). Altogether, during embryonic development, we propose that DFPs must undergo an intrinsic reprogramming process to commit to fibroblast lineages and differentiate toward the terminal fates.

Epigenetic regulators have been extensively studied in the context of cellular differentiation (Boyer et al., 2006; Flora and Ezhkova, 2020; Kang et al., 2019; Lee et al., 2006; Plikus et al., 2021). A proposed mechanism of chromatin regulation is through the addition of me3 to H3K27, which is associated with condensed and silenced chromatin while removal of me3 is associated with accessible and active chromatin (Agger et al., 2007; Cao et al., 2002; Kang et al., 2019; Xiang et al., 2007). The most widely known regulators of methylation on H3K27 are part of the Polycomb repressive complex 2 (PRC2), EZH1 and EZH2. In the epidermis, the addition and removal of the trimethyl group (me3) on histones-H3 Lysine 27 (H3K27) were shown to be critical in controlling the differentiation process of epidermal stem cells during development (Ezhkova et al., 2009; Sen et al., 2008). Demethylases, such as Kdm6a/Utx and Kdm6b/Jmjd3, specifically remove me3 and de-repress chromatin (Agger et al., 2007; Xiang et al., 2007). In dermal fibroblasts, EZH2 has been shown to act as an epigenetic rheostat to control the Wnt and Retinoic Acid signaling in DFPs (Thulabandu et al., 2021). However, a full understanding of how the methylation status of H3K27 regulates fibroblast differentiation has not been shown. In addition, other studies have also investigated the roles of different epigenetic marks in regulating and maintaining fibroblast lineages in developed skin. Acetylation of H3K27 and HDAC2 activities are required for the maintenance of Papillary Fibroblast lineage postnatal (Kim et al., 2022). H3K4me3, H3K9me3, and H3K27me3 all display reduced methylation levels in the hair follicle stem cells prior to hair growth, and inhibition of hypomethylation delays skin wound healing(Kang et al., 2020; Lee et al., 2016). Nonetheless, an epigenetic regulation that programs dermal fibroblast lineage commitment and differentiation during embryonic development remains unknown.

The recent emergence of single-cell technology has advanced the study of fibroblast heterogeneity drastically (Plikus et al., 2017). The transcriptomic changes in restricted fibroblast lineages, specifically in the hair follicle-associated fibroblasts, have been characterized at different timepoints that regulate Dermal Papilla establishment (Biggs et al., 2018; Gupta et al., 2019; Mok et al., 2019). In our recent study of transcriptomic and chromatin landscapes at the single-cell level, we showed that the transcriptomic profiles of fibroblast lineages could only reflect the cellular states, while the accessible chromatin profiles can predict the lineage restriction and terminal fate commitment (Thompson et al., 2022). Overall, we hypothesize that the differentiation of dermal fibroblast progenitors could be defined by incorporating chromatin accessibility profiles and epigenetic regulation.

To establish the multimodal reference point for dermal fibroblast progenitors, we compared the molecular states of dermal fibroblast progenitors (DFP) at E14.5 to late embryonic fibroblasts at E18.5 using single-cell transcriptomics, single-cell ATAC, ChIP-seq, genetic ablation of epigenetic regulator Jmjd3/Kdm6b, and functional chamber allografting assays. We found that DFPs at E14.5 are predicted to differentiate into hair follicle-associated fibroblast, but they are not intrinsically programmed as they failed to support hair follicle formation *ex vivo.* E14.5 DFPs possess a closed-off chromatin profile, marked by a high level of H3K27me3, and they require epigenetic programming mediated by Kdm6b/Jmjd3 to commit to specific fibroblast lineages. Finally, we share our processed data for all our experiments on our website skinregeneration.org.

## RESULTS

### DFPs differentiate into distinct fibroblast lineages in vivo but fail to regenerate functional skin in grafting assays

Using lineage tracing assays, we have previously shown that DFPs are present throughout the dermis at E14.5 and that a single DFP has the potential to contribute to the formation of all layers and fibroblast subtypes (Driskell et al., 2013; Rognoni et al., 2016). We hypothesized that E14.5 fibroblasts are multipotent DFP progenitors in developing murine skin. We performed scRNA-seq from whole skin samples of E14.5, E17.5, and P5 mice (Figure 1A-E). Altogether, we captured and sequenced a total of 43,740 cells, of which 17,356 cells were classified as dermal fibroblasts (Figure S1A-B). To evaluate DFP differentiation and developmental potential, we integrated and batch-corrected all three time points together and embedded the single-cell clusters as a PAGA map to improve interpretation but also preserve the global topology of cells transition (Figure 1D-E) (Hie et al., 2019; Wolf et al., 2019). DFPs at E14.5 were classified into two distinct subsets based on their horizontal anatomical positions as previously published (Gupta et al., 2019; Jacob et al., 2022). Upper DFPs (clusters 0, 2,11) expressing Crabp1, Cav1, Nkd1, and Lef1 resided closer to the epidermis, while Lower DFPs expressing Thbs1, Ptn, and Mfap5 (clusters 1,5,6) were located near the panniculus carnosus (Figure 1A, D-E, Figure S1C). Furthermore, fibroblast subsets in E17.5 and P5 populations were identified based on gene expression profiles that have been previously defined (Figure S1) (Guerrero-Juarez et al., 2019; Phan et al., 2020; Thompson et al., 2022). We identified clusters of Dermal Papilla, Papillary Fibroblasts, Reticular Fibroblasts, Fascia, and Pre-adipocytes in both E18.5 and P5 skin. In addition, clusters of mature papillary and ECM-producing fibroblasts were detected only in P5 skin, reflecting the maturation of dermal fibroblasts post-natal. Our PAGA integrated analysis revealed that most E17.5 fibroblasts were closely associated with E14.5 DFPs while P5 fibroblasts clustered distinctly based on their lineage functions, except in the lineage-committed specialized fibroblasts such as Dermal Papilla and Pre-adipocytes, where E17.5 and P5 clustered together (Figure 1D-E).

**Figure 1.**
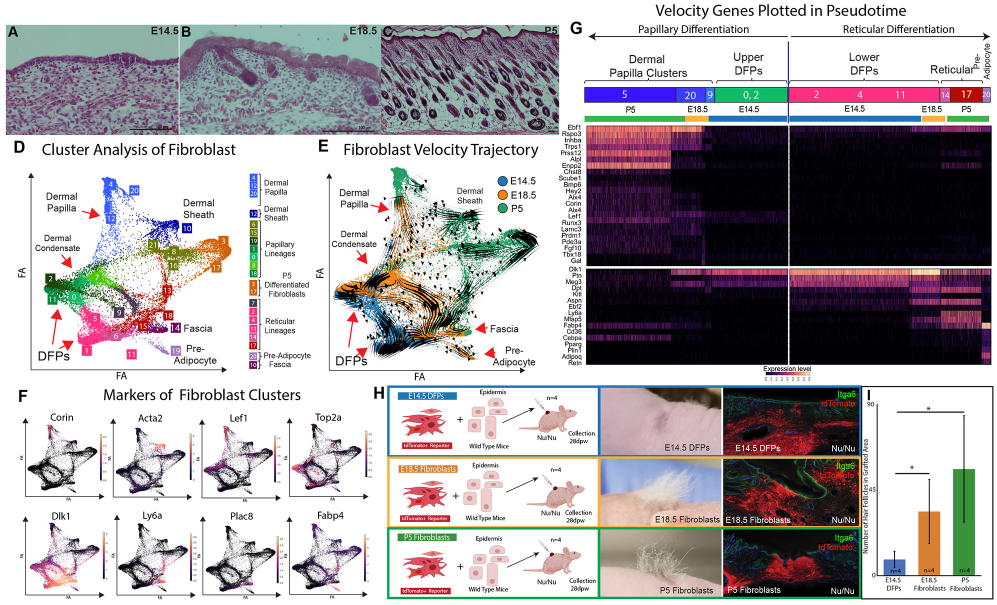
DFPs are not intrinsically programmed to support skin reformation. (A-C) H&E histology of murine skin sections at Embryonic Day E14.5 (A), E17.5 (B), and Post-natal day 5 (P5) (C). (D-E) PAGA-initialized single-cell embedding of fibroblasts of 3 timepoints integrated. (F) Expression of Fibroblast Marker Genes. (G) Pseudotime-heatmap depicting the differentiation projections of Upper Progenitors to Dermal Papilla and Lower Progenitors to Pre-adipocyte. (H-J) *Ex vivo* chamber grafting assay testing the ability of fibroblast populations at E14.5, E17.5, and P5 to support hair follicle formation. (J) Histology of reconstituted skin in chamber grafting assay. (J) Quantification of the number of hair formations in grafted area.

To further establish the continuous transition between DFPs and fibroblast lineages, we performed RNA velocity analysis on the PAGA-embedded fibroblast subsets, which accounted for the transcriptional dynamics of splicing kinetics within each cell (Bergen et al., 2020). Upper DFPs are shown to differentiate into papillary fibroblasts (clusters 7, 8, 16, 21), dermal papillae (clusters 4, 12, 20), dermal sheath (clusters 10), and mature fibroblasts (clusters 3, 17), while lower DFPs are projected to become Upper DFPs, reticular (clusters 9, 13, 15, 18), fascia (cluster 14) and pre-adipocytes (clusters 19) (Figure 1D-E) (Correa-Gallegos et al., 2019; Driskell et al., 2014; Guerrero-Juarez et al., 2019; Joost et al., 2020; Shook et al., 2020). RNA velocity analysis also provided us with a list of putative driver genes that need to be activated for cells to differentiate toward a specific cluster. Using these genes, we generated a heatmap depicting the pseudotime transitions of Upper and Lower DFPs into two distinct lineages: Dermal Papilla and Pre-adipocytes (Figure 1G). We found that multiple velocity driver genes of E17.5 and P5 Dermal Papilla were expressed as early as E14.5 Upper DFPs (Figure 1G). Overall, our results suggested that at E14.5, DPFs have the potential to differentiate into specialized cell types to support and maintain the different dermal niches in post-natal skin.

Grafting assays, in which cells are transplanted into host tissues and allowed to interact with the surrounding microenvironment, provide a powerful tool to assess the differentiation potential and lineage restrictions of stem cells *ex vivo* (Driskell et al., 2013; Jensen et al., 2010). To functionally challenge the potential of E14.5 DFPs to support the formation of fully functional skin, we performed a skin reconstitution experiment utilizing the chamber grafting technique. We collected whole, unsorted DFPs from E14.5 and compared their ability to reform hair follicles and skin from fibroblasts from E17.5 and P5 tdTomato-positive mice to trace their contribution within grafted tissue. A combination of dermal cells and epidermal cells were injected into a silicone chamber grafted on top of an immunocompromised Foxn1^-/-^ mouse (Figure 1K). Skin from areas of the graft were collected at 28 days post-surgery to assess the reformation and development of skin. The analysis and quantification of the grafting assay revealed that E17.5 and P5 differentiated fibroblast populations were able to support the reformation of fully functional hair follicles and hair fibers while E14.5 DFPs failed to regenerate and develop hair follicles (Figure 1L). This result suggests that, despite the potential trajectory and presence of early dermal papilla cells, DFPs from E14.5 skin are not intrinsically programmed to differentiate ex vivo to support fully functional skin reformation.

### DFP programming occurs by specifically increasing the accessibility of chromatin architecture in distinct lineages

The chromatin accessibility landscape regulates cellular gene expression and is highly informative of its states and fates of cells (Buenrostro et al., 2015, 2013; Cao et al., 2017; Cusanovich et al., 2018, 2015; Thompson et al., 2022; Trapnell, 2015). We hypothesized that the intrinsic differentiation program within E14.5 DFPs could be reflected in its chromatin accessibility profiles. We performed single-cell ATAC-seq assay on murine DFPs at E14.5 and differentiating fibroblasts at E18.5. After quality control, filtering, and pre-processing, 10,913 fibroblasts from both time points were used for downstream analyses including dimension reduction and unsupervised clustering (4,187 from E14.5 and 6,006 from E18.5) (Figure 2A-B). Standard quantification of the ATAC peak number per cell comparing E14.5 and E18.5 fibroblasts revealed a large increase in open chromatin regions during the differentiation process (Figure 2C). Differential peaks analysis was performed using Wilcoxon testing between Time identified 6200 peaks in E14.5 and 13566 peaks in E18.5 fibroblasts. Coverage plots depicting the significant changes in open chromatin regions between E14.5 and E18.5 highlighted the increase of accessible peaks during the differentiation process of DFPs (Figure 2D). Our results indicated that chromatin reprogramming of DFPs during differentiation is associated with a selective, de-repressive, and opening of the genome.

**Figure 2.**
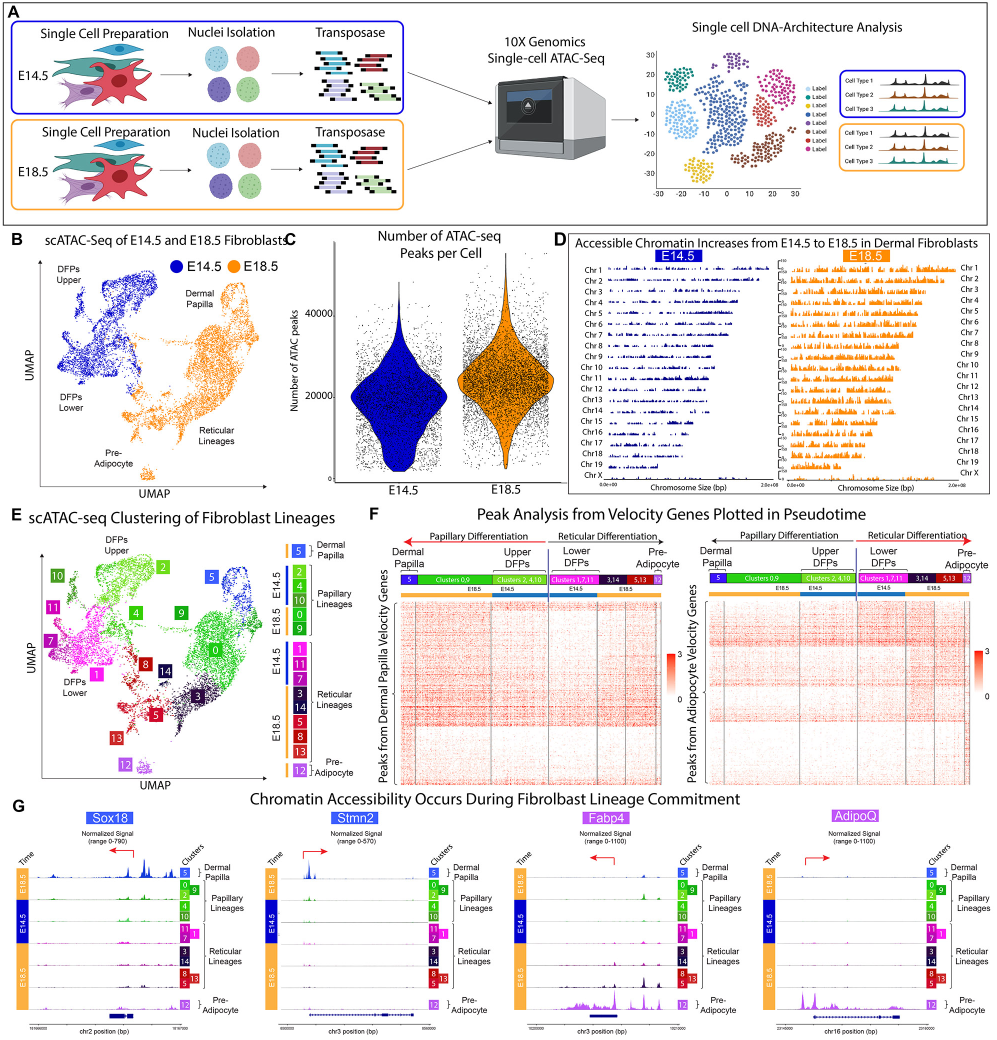
DFP differentiation is defined by inaccessible chromatin architecture. (A) Experimental design of scATAC-seq experiments comparing E14.5 and E18.5 dermal fibroblasts. (B) UMAP plot showing the distinct clustering of E14.5 and E18.5 fibroblasts. (C) Coverage Plots highlighting the significant increase in open accessible chromatin regions from E14.5 to E18.5 fibroblasts. (D) Quantification of Total ATAC Peaks per cell. (E) UMAP plot showing the classification of fibroblast cell types in scATAC-seq data. (F) Pseudotime-heatmap of scATAC-seq peaks associated with RNA velocity genes required for Dermal Papilla and Pre-adipocyte differentiation. Dermal Papilla and Pre-adipocyte populations are flanking the E14.5 DFPs which are located in the centrally (G) Tracks-plot arranged in pseudotime showing specific accessibility regions of Sox18 and Stmn2 in Dermal Papilla, Fabp4 and AdipoQ in Pre-adipocytes.

To investigate the relationship between the de-repression of chromatin and the establishment of fibroblast heterogeneity from DFPs, unsupervised clustering of scATAC-seq data was performed resulting in 15 distinct clusters from both time points (Figure 2E, S2A). Cross-modality integration and label transfer were used to integrate scATAC-seq with scRNA-seq data for cluster identification (Methods, Figure S2B) (Stuart et al., 2021). Consistent with our transcriptomic analysis, DFPs at E14 were split into Upper DFPs and Lower DFPs, while the E18 fibroblasts contained Dermal Papilla, Papillary, Reticular, and Pre-adipocytes clusters (Figure 2E). Interestingly, even though Dermal Papilla were detected histological and in our scRNA analysis, we did not detect them in our scATAC-seq data (Figure 2E). Differential peak analysis revealed distinct chromatin landscapes among fibroblasts at E18.5, while the E14.5 Upper and Lower DFPs possessed less unique accessible peaks profiles, which was consistent with the opening of chromatin during DFPs differentiation processes (Figure S1D). Similar to P0 fibroblasts(Thompson et al., 2022), E18.5 fibroblasts also shared a gradient of significant peaks within their respective lineages, with the differentiated Dermal Papilla and Pre-adipocytes fates possessing the most distinct chromatin profiles (Figure 2G, S2D-E). Considering the predicted RNA velocity trajectories of DFPs to differentiate into Dermal Papilla and Pre-adipocyte, we hypothesized that the chromatin accessibility profiles of the equivalent cell types will also reflect a similar differentiation trajectory. We constructed a pseudotime heatmap based on our scATAC-seq data but defined by RNA velocity driver genes for each fibroblast cell type from DFPs to Papillary to Dermal Papilla and DFPs to Reticular to Pre-adipocyte (Methods, Figure 2F). We found that as Upper DFPs differentiated toward Papillary and Dermal Papilla, the velocity driver genes of Papillary and Dermal Papilla became more accessible (Figure 2F). Notably, the driver genes of Upper DFPs were accessible across all fibroblast subtypes. Dermal Papilla had accessible chromatin regions for Papillary driver genes, but DP-specific genes were not accessible in Papillary, indicating the differentiated fate of Dermal Papilla as previously reported (Figure 2F). This finding confirmed the continuous transition of Upper DFPs to Papillary to Dermal Papilla, where Upper DFPs require making their chromatin landscape specifically accessible during their differentiation process. The closed chromatin profiles of DFPs at E14.5 without accessible regions for Dermal Papilla driver genes might help explain the inability of DFPs to reform hair in the chamber grafting assays. Similarly, the Lower DFPs differentiation trajectory was also observed using Lower DFPs, Reticular, and Pre-adipocytes RNA velocity driver genes (Figure 2F). Overall, our analysis revealed that during differentiation, DFPs specifically make their chromatin accessible in regions associated with the specific driver genes at E18.5 in a stepwise manner (Figure 2F). We also concluded that the differentiated fibroblast fates of Dermal Papilla and Pre-adipocytes required unique reprogramming of dermal fibroblasts with specifically accessible regions such as Sox18, Stmn2 for Dermal Papilla and Fabp4, AdipoQ for Pre-adipocytes (Figure 2G).

### Chromatin programming of DFPs is associated by the removal of the repressive histone modification H3K27me3

Post-translational modifications of histones can regulate gene accessibility and expression during the differentiation process in various tissue models (Boyer et al., 2006; Heintzman et al., 2009; Karlić et al., 2010; Lee et al., 2006). Specifically, the Histone 3 Lysine 27 trimethylation mark (H3K27me3), which is associated with the repression of gene transcription, plays a key role in the differentiation process of epidermal keratinocytes (Ezhkova et al., 2009; Kang et al., 2019; Sen et al., 2008). Our scATAC-seq data suggested that the opening of chromatin regions at lineage-specific genes was critical for fibroblast differentiation. To study the role of H3K27me3 in regulating open chromatin regions of DFPs during embryonic differentiation, we performed H3K27me3 ChIP-seq comparing E14.5 and E18.5 Fibroblasts (Figure 3A). In conformity with the increase in open chromatin regions between E14.5 and E18.5 fibroblasts, we found a sharp decrease of 52% in the number of H3K27me3 peaks (from 19060 peaks at E14.5 to 8768 peaks at E18.5, n=2) by visualizing the heatmaps and coverage plots of peaks called at each time point (Figure 3B-D). In addition, we also confirmed that the majority of the genes associated with H3K27me3 peaks at E18.5 were presented at E14.5 (95%), indicating that there was a significant reduction in H3K27me3 associated genes and not a dynamic change where undifferentiated genes were repressed and differentiated genes were de-repressed (Figure 3E). We next utilized DiffBind to identify the significant changes between E14.5 and E18.5 H3K27me3 peaks in DFs, which resulted in 4699 significant peaks that decreased from E14.5 to E18.5 and only 29 increase peaks (Figure 3F). To investigate their functions, Gene Ontology analysis of genes associated with the significant de-repressed peaks from E14.5 revealed the biological processes associated with skin differentiation and development such as multicellular organism development, cell differentiation, neuron differentiation, Canonical Wnt signaling pathway, hair follicle development, and glucose homeostasis (Figure 3G). Within this list, we specifically found genes that played important roles in the establishment and differentiation of dermal niches such as Sox18 and Alpl for Dermal Papilla and Cebpa and Pparg for Pre-adipocytes (Figure 3H). Overall, our data revealed that the changes in chromatin landscapes in DFPs between E14 and E18 coincided with the demethylase of H3K27me3 markers within genes required for DFPs differentiation. More importantly, we also found an active, high level of expression for the H3K27me3 specific demethylase – Kdm6b/Jmjd3 – in both E14 and E18 fibroblasts (Figure 3I).

**Figure 3.**
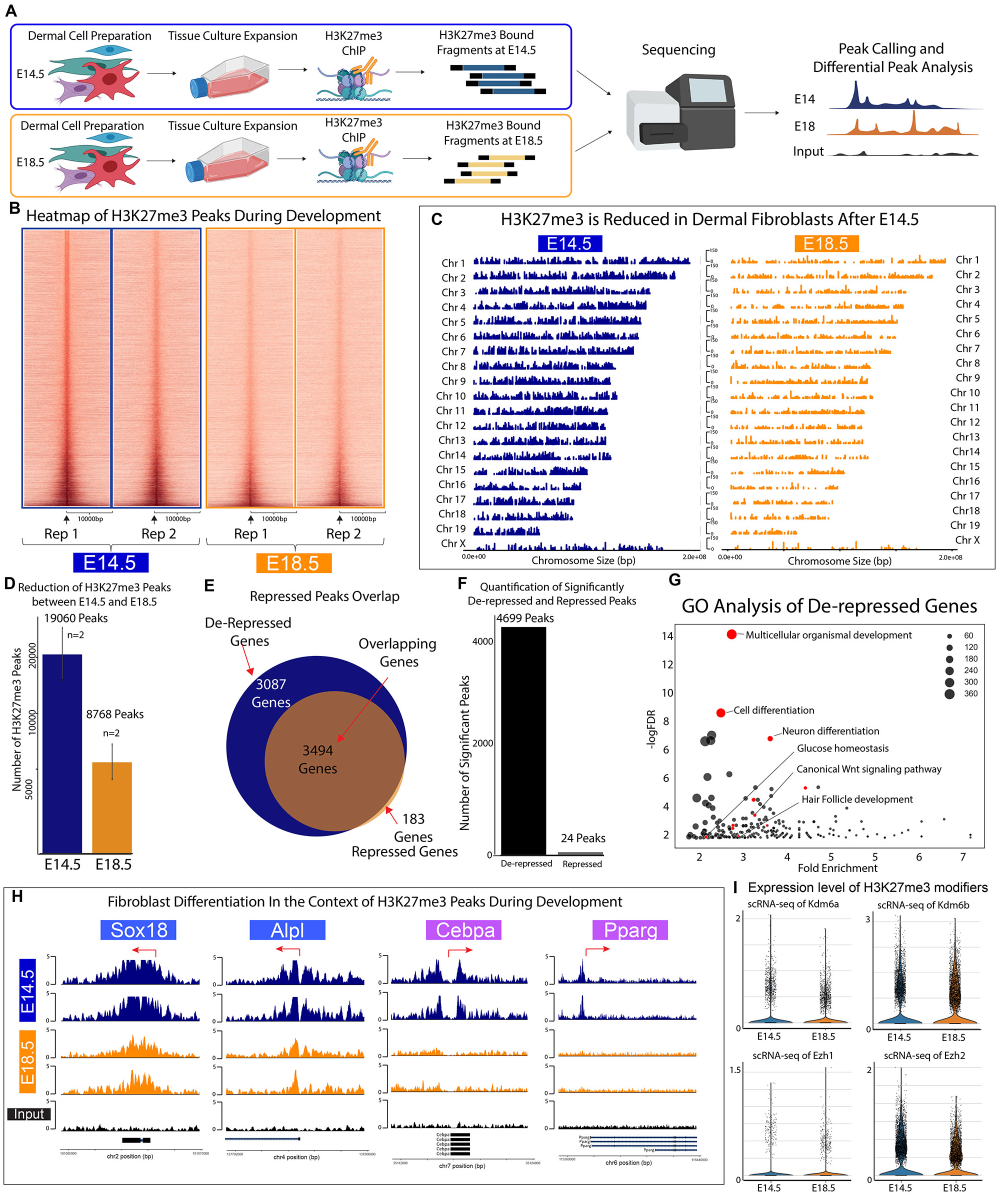
Differentiation of DFPs during embryonic development is coupled with demethylation of H3K27me3. (A) Experimental design of H3K27me3 ChIP-seq comparing E14.5 and E18.5 Fibroblasts. (B) Heatmaps of H3K27me23 Peaks comparing E14.5 and E18.5 Fibroblasts. (C) Coverage Plots highlighting the reduction of H3K27me3 from E14.5 to E18.5 dermal fibroblasts. (D) Quantification showing the decrease in H3K27me3 peaks (19060 peaks at E14.5 and 8768 peaks at E18.5, n=2) (E) Venn diagram revealing the distribution of genes associated with H3K27me3 peaks changes between fibroblasts from E14.5 to E18.5. (F) Quantification of significantly differentiate H3K27me3 binding between E14.5 and E18.5 fibroblasts (4699 peaks at E14.5 and 24 peaks at E18.5) (G) GO Analysis of Significantly De-repressed Genes revealed important Fibroblast differentiation processes. (H) Tracksplot of important de-repressed genes associated with fibroblast differentiation (Sox18 and Alpl for Dermal Papilla, Cebpa and Pparg for Pre-adipocyte). (I) scRNA-seq showing the expression level of H3K27me3 demethylase Kdm6a and Kdm6b, methyltransferase Ezh1 and Ezh2 at E14.5 and E18.5.

### Genetic ablation of Histone Demethylase Kdm6b/Jmjd3 inhibits DFPs differentiation

Previous studies have identified the roles of Ezh2 and PRC2 complex in controlling epidermal differentiation by adding the H3K27me3 marker at specific genes to maintain the stemness of the progenitor cells. In contrast, the H3K27me3 demethylase Kdm6b/Jmjd3 has also been shown to be critical in epidermal stem cell differentiation, where ablation of Jmjd3 leads to inhibited differentiation of this cell type (Ezhkova et al., 2009; Sen et al., 2008). Based on the significant decrease in H3K27me3 markers of DFPs from E14 to E18, we hypothesize that Kdm6b/Jmjd3 demethylase activity is required for the differentiation of dermal fibroblasts. To test the role of Kdm6b/Jmdj3 in dermal differentiation, we generated a transgenic dermal-specific knockout of Jmjd3 mouse model driven by activation of Dermo1 in dermal fibroblasts (Figure 4A). The Dermo1Cre-Kdm6b^fl/fl^ (Kdm6b-cKO) mice are neonatally lethal, and we focused our study on the changes occur between E14.5 and E18.5. Histological analysis of E14.5 skin showed similar stages of skin development between Kdm6b-cKO skin and the WT control where a thin layer of the dermis was found in conjunction with the formation of dermal condensates (Figure 4B). However, at E18.5, we observed drastic changes as Kdm6b-cKO skin appeared to be underdeveloped with smaller hair follicles and significantly thinner dermal layers (Figure 4C-D). We also found that the development of one differentiated cell type – Dermal Papilla – was also inhibited, with fewer Dermal Papilla per hair follicle (Figure 4E). Flow cytometry staining for CD133/Prom1, a surface marker of Dermal Papilla, confirmed a significant 20 % decrease in their number from whole dermal preparation (Figure 4F).

**Figure 4.**
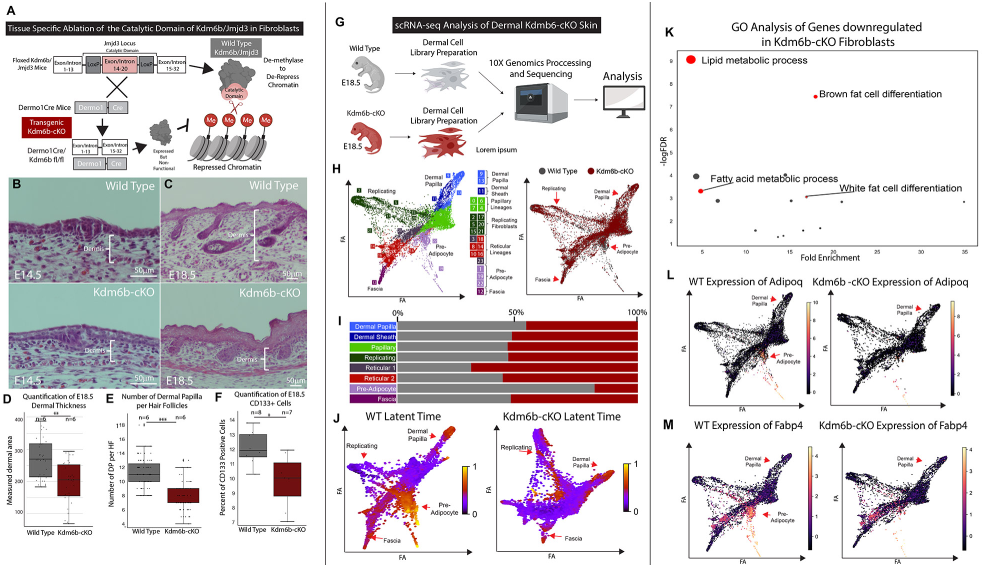
Kdm6b/Jmjd3 regulates DFP differentiation. (A) Transgenic approach to specifically knock-out Jmjd3 in murine model. (B-C) H&E staining for histological analysis of E14.5 and E18.5 WT and Kdm6bKO skin. (D) Quantification of E18.5 thickness comparing E18.5 WT and Kdm6bKO skin (p-value <0.01) (E) Changes in number of Dermal Papilla per Hair Follicle between WT and Kdm6bKO. (F) Quantification of flow cytometry comparing the percentages of CD133 positive cells between E18.5 WT and Kdm6bKO dermal prep. (G) Experimental strategy to generate single-cell RNA-seq data from E18.5 WT and Kdm6bKO mice. (H) PAGA embedded plots showing cell clusters of integrated E18.5 WT and Kdm6bKO fibroblasts. (I) Bar plots showing the contribution of each condition (WT and Kdm6bKO) to each fibroblast sub-cell types. (J) Comparison of RNA velocity predicted latent time between E18.5 WT and Kdm6bKO fibroblasts. (K) GO analysis of Genes significantly downregulated in Kdm6bKO fibroblasts. (L) PAGA plots showing the expression levels of Adipogenic genes AdipoQ and Fabp4 between WT and Kdm6b KO.

To extensively characterize the changes induced by Jmjd3 ablation, we performed a scRNA-seq experiment comparing the E18.5 dermis of Kdm6b-cKO mice to the littermate genetic control (Figure 4G). After pre-processing and filtering steps, 12,947 fibroblasts were retrieved for downstream analysis (Figure 4H). PAGA embedding of single cell clusters were performed to visualize the trajectory and continuous transition among different fibroblast cell types. In addition, we performed cluster classification as in Figure 1D-F (Figure 4H, Figure S2C). Interestingly, we did not observe a significant change in the number of Dermal Papilla between WT (Kdm6b^fl/fl^) and Kdm6b-cKO. Instead, we found that there was a dramatic reduction in number of pre-adipocytes detected in Kdm6b-cKO skin compared to the WT control (1081 in WT, 238 in Kdm6b-cKO), along with a large decrease in the number of lymphatic vasculatures (Figure 4H-I, Figure S2A-B). This observation coincides with previous studies reporting the relationship between dermal fibroblasts, dermal pre-adipocytes and endothelial vasculature niche (François et al., 2008; Gupta et al., 2012; Jiang et al., 2014; Martinez-Corral et al., 2015; Oliver et al., 2020; Pichol-Thievend et al., 2018; Stone and Stainier, 2019; Tang et al., 2008). To further interrogate the effect of Kdm6b ablation, we performed the RNA velocity and latent time analysis comparing WT to Kdm6b-cKO fibroblasts. Latent time analysis represents the cellular internal clock and approximation of cell state as they differentiate using only the intrinsic transcriptional dynamics (Bergen et al., 2020). The analysis revealed changes in the terminally differentiated cell types where Kdm6b-cKO fibroblasts shifted differentiated cell types from Pre-adipocytes to Replicating fibroblasts (Figure 4J). We next performed a differentially expressed genes analysis compared the two conditions using Diffxpy, a more optimized package for statistical testing of single-cell data (Methods). We found that there are 472 genes downregulated in Kdm6b-cKO fibroblasts compared to WT. GO analysis of this list genes revealed the difference in genes expression were related to the formation and differentiation of the Pre-adipocytes population, where biological processes such as Lipid metabolic process, Brown fat cell differentiation, Fatty acid metabolic process, and White fat cell differentiation were significantly enriched. These differences were driven by the significant changes in the number of Pre-adipocytes between the two conditions, so genes such as Fabp4, Adipoq are significantly downregulated in Kdm6b-cKO fibroblasts (Figure 4L-M). In summary, our histological and scRNA-seq analyses revealed that the ablation of Kdm6b/Jmdj3 demethylase activity in the dermis inhibited the differentiation of DFPs to specific fibroblasts cell fates of Dermal Papilla and Pre-adipocytes, while the transient states of Papillary and Reticular were less impacted.

### Kdm6b regulates the epigenetic programming of DFP differentiation into functional Dermal Papilla

Despite the significant impact on the development of the dermis and Dermal Papilla, scRNA-seq data of Kdm6b-cKO skin did not capture large changes in genes associated with dermal papilla function. We hypothesize that the dermal papilla transcriptomic profile reflects the specialized niche environment and epidermal signals, and that Kdm6b/Jmjd3 ablation will alter the intrinsic H3K27me3 epigenetic programs of Kdm6b-cKO fibroblasts. To investigate the altered epigenetic profile, we performed H3K27me3 ChIP-seq comparing E18.5 WT and Kdm6b-cKO fibroblasts (Figure 5A). We found that E18.5 Kdm6b-cKO fibroblasts have an increase in the number and level of H3K27me3 peaks (Figure 5 B-D). Compared to the average of 26,228 peaks called in E18.5 WT, Kdm6b-cKO fibroblasts had a large increase of peaks to 42331 H3K27me3 peaks (Figure 5B). Visualizations of H3K27me3 peaks using heatmaps (Figure 5C) and coverage plots across the whole genome (Figure 5D) highlighted the apparent increase in H3K27me3 repressive peaks of the same regions in Kdm6b-cKO fibroblasts, since the majority of called peaks overlapped between WT and Kdm6b-cKO (Figure S3A). We next performed differential peaks analysis using DiffBind to identify significant changes between the two conditions, which resulted in 1916 Kdm6b-cKO peaks compared to 29 peaks in WT fibroblasts (Figure S3B). GO analysis of the Kdm6b-cKO revealed significantly increased peaks for enrichment of biological processes such as Potassium ion transmembrane transport, Signal transduction, Cell differentiation, Multicellular organism development, Cell-cell signaling, and Neuron differentiation (Figure S5C). We did not observe processes directly related to skin, hair follicle, and dermal white adipocyte differentiation being enriched, even in the present of important genes for Dermal Papilla and Pre-adipocyte like Sox18, Cebpa, Hes5, Rspo1, and Eya2 (Figure 5E) (Mok et al., 2019; Sennett et al., 2015). When comparing with the transcriptomic changes induced by Kdm6b ablation, we found an overlap of 38 genes between genes downregulated in Kdm6b-cKO scRNA-seq and increase in ChIP-seq data, with many of them have been shown to be important in the development of Dermal Papilla (Figure S3D). Our results confirmed that genes associated with dermal papilla function are regulated by Kdm6b/Jmjd3 de-methylase at the epigenetic level, while other epigenetic modulators could compensate for Jmdj3 function.

**Figure 5.**
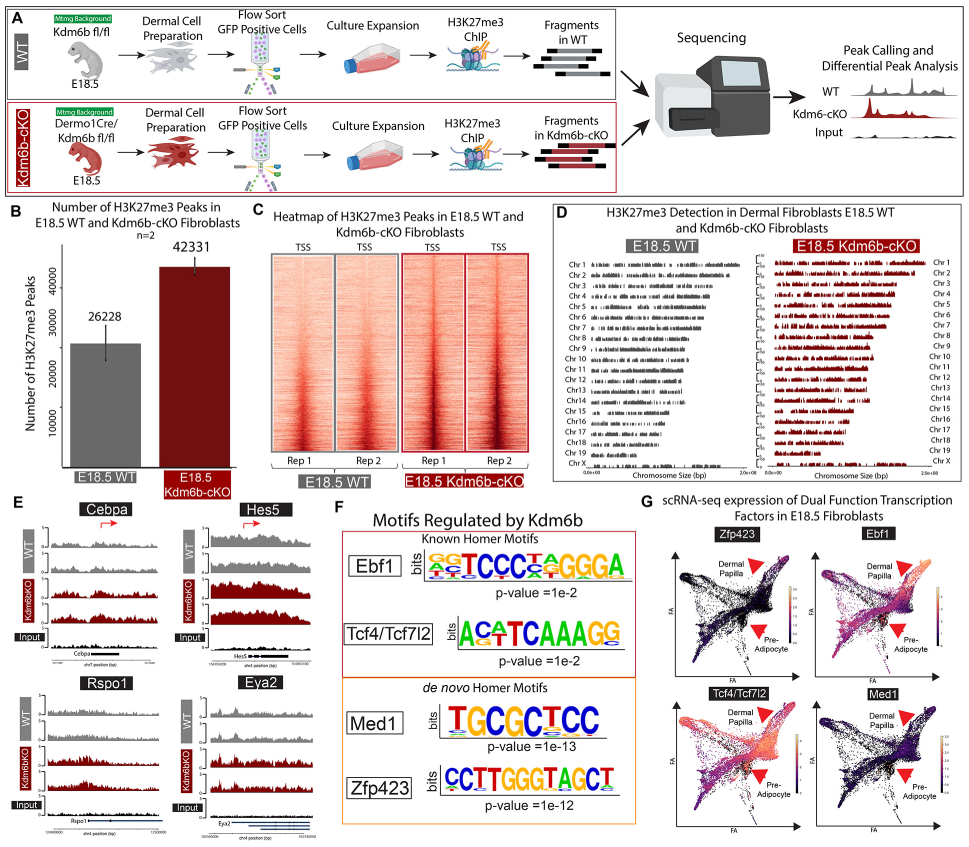
Kdm6b regulates epigenetic de-repression of dual-function Adipogenic and Dermal Papilla transcription factors. (A) Experiment design for H3K27me3 ChIP-seq comparing E18.5 WT and Kdm6bKO. (B) Heatmaps depicting the changes in H3K27me3 peaks in Kdm6bKO fibroblasts. (C) Coverage plots highlighting the increase in H3K27me3 peaks in Kdm6bKO. (D) Bar plots showing the number of H3K27me3 peaks called in E18.5 WT and Kdm6bKO Fibroblasts. (E) Tracksplots of relevant transcription factors loci for Dermal Papilla and Pre-adipocyte lineage commitment (Cebpa, Prdm16, Bmp6, and Hes5). (F) Homer motif analysis of significantly repressed peaks in Kdm6bKO fibroblasts. (G) scRNA-seq expression levels of Motifs regulated by Kdm6b activity.

To assess the role of epigenetic regulation in Dermal Papilla functions, we hypothesized that the ablation of Kdm6b/Jmjd3 in DFPs will lead to inaccessibility of transcription factor binding sites that regulate fibroblast lineage functions. To address this hypothesis we utilized the motif finder Homer to screen for binding motif enrichment in the significant peaks regulated by Kdm6b (Figure 5F)(Heinz et al., 2010). The known Homer motifs revealed an enrichment of Ebf1, Tcf4/Tcf7l2 binding motifs in peaks that were increased in Kdm6b-cKO fibroblasts, while the *de novo* Homer motifs detected the significant enrichment of dermal adipocyte differentiation transcription factors Med1 and Zfp423 (Figure 5F) (Biferali et al., 2021; Gupta et al., 2012, 2010; Harms et al., 2015; Shao et al., 2016). Interestingly, the expression of these transcription factors was not specific to just Dermal Papilla or Pre-adipocyte but were expressed throughout fibroblasts populations (Figure 5G). This result suggested that they might have distinct roles in both Adipogenesis and Dermal Papilla functions.

Next, we wanted to investigate how Kdm6b/Jmjd3 might influence Dermal Papilla function, in the context of these dual-function transcription factors such as Ebf1 and Zfp423. Early B cell factor 1 (Ebf1) has been shown to regulate the formation of dermal white adipocytes and hair follicle cycles (Festa et al., 2011). Consequently, we hypothesized that the differential binding sites for these dual-function transcription factors are regulated by Kdmb6b/Jmjd3 (Figure 6A). We performed a motif matching analysis of Ebf1 motif using scATAC peaks specific to Dermal Papilla and Pre-adipocytes, which revealed a distinct but non-exclusive Ebf1 regulomes in both cell types with 1724 shared genes, 1988 genes specific to Dermal Papilla, 2101 genes specific to Pre-adipocytes (Figure 6B, Figure S5). We found Ebf1 motifs in Sox18, Lamc3, and Fgf10, regulatory regions that were specifically accessible in Dermal Papilla cells (Figure 6C). GO analysis of genes that possessed accessible Ebf1 binding motif in Dermal Papilla included biological processes such as Multicellular organism development, Negative regulation of cell proliferation, Angiogenesis, Wnt signaling pathway, and Hair follicle development. The counterpart in Pre-adipocytes involved Protein phosphorylation, apoptotic process, negative regulation of Wnt signaling pathway, Response to hyperoxia, and Lipid metabolic process (Figure 6B). This result demonstrated the dual roles of Ebf1 in two distinct lineages where a single transcription factor can regulate unique subset of genes to exert diverged functions, and Kdm6b/Jmdj3 mediates specific transcription factor functions by de-repressing different genes regions within each lineage. To confirm that Kdm6b-mediated epigenetic program is required for the intrinsic reprogramming of DFPs to Dermal Papilla, we performed another chamber allografting assays comparing E18.5 dermis of WT and Kdm6b-cKO mice (Figure 6D). We found that Kdm6b-cKO dermis could not support fully functional skin in grafting assay by the lack of hair follicles and hair fibers reformations, similar to E14.5 WT dermis (Figure 1H), while E18.5 WT dermis can reform numerous functional hair follicles and hair fibers (Figure 6D). Overall, we concluded that DFPs differentiation into distinct cell fates and lineages were programmed by the establishment of unique open chromatin profiles, which is mediated by Kdm6b demethylase activity removing H3K27me3 markers as specific, functional genes loci.

**Figure 6.**
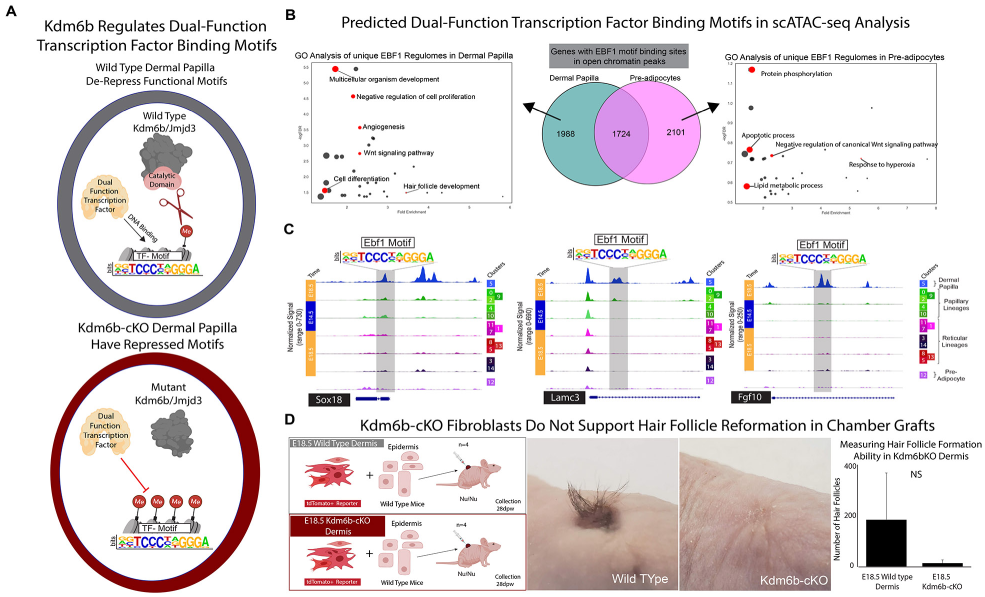
Kdm6b regulates DFP differentiation into fully functional Dermal Papilla. (A) Model for Dual-Function Transcription Factor regulation of Dermal Papilla activity regulated by Kdm6b/Jmjd3 de-repression. (B) Dual-Function Transcription Factor, Ebf1, binding activities represented by a GO-analysis of Ebf1 motifs in scATAC-seq peaks from Dermal Papilla and Pre-Adipocytes fibroblast populations. (C) scATAC-seq track-plot of Dermal Papilla genes, Sox18, Lamc3, and Fgf10. Predicted Ebf1 motifs are highlighted in a grey box on top of the peak that is predicted to have the motif. (D) Experimental strategy for skin reconstitution assay (chamber grafting assay) comparing E18.5 Fibroblast of WT and Kdm6bKO skin. Images of hair follicle formations within grafted area comparing WT and Kdm6bKO Fibroblasts and quantification of hair follicles reformed in chamber grafting assays.

## DISCUSSION

### Defining the epigenetic state of a dermal fibroblast

Recent studies have demonstrated that stem cells require extensive chromatin accessibility priming prior to lineage commitment (Buenrostro et al., 2018; Ranzoni et al., 2021). The accessible chromatin profiles of dermal fibroblasts and other cell types could reflect their potential to differentiate into a terminal fate (Adam et al., 2018; Kim et al., 2022; Li et al., 2017; Ma et al., 2020; Thompson et al., 2022). Here, we discovered that DFPs possess inaccessible chromatin profiles at differentiation specific genes required for lineage commitment. The inaccessible and unprogrammed chromatin profiles of E14.5 DFPs compared to the differentiating fibroblasts helped explain the inability of E14.5 dermis to support HF reformation in the chamber grafting assay (Figure 1, Figure 2D). Moreover, our scATAC-seq data confirmed the bifurcation of distinct fibroblast lineages during embryonic development (Driskell et al., 2013; Sorrell and Caplan, 2004). We found that Upper DFPs possessed the most stem-like chromatin profiles with the least number of accessible peaks. Papillary and Dermal Papilla are closely associated with each other, where significant differential peak analysis and integration with RNA velocity driver genes showed the large overlapping regions between them (Figure 2F, S1D). In addition, our integrative RNA velocity and scATAC-seq analysis revealed a stepwise arrangement of accessible chromatin for DFP differentiation, where Dermal Papilla possess the most unique chromatin architecture, but genes specific for Papillary fibroblasts also remain accessible in the Dermal Papilla (Figure 2F). This result confirmed our previous finding of fibroblasts states and fates, with the Dermal Papilla and Pre-adipocytes being two defined terminal fates for DFP differentiation (Thompson et al., 2022).

Our investigation of the regulation of chromatin architecture by epigenetic histone modifications revealed that DFP differentiation occurs through specific de-repression of chromatin to support fibroblast lineage function. We found that DFPs required a significant decrease in H3K27me3 levels in genes associated with differentiating fibroblasts functions such as Sox18, Alpl, Cebpa, Pparg, etc. Notably, during the differentiation process, we only observed a small increase in H3K27me3 markers (24 peaks) (Figure 3B-F). Our result suggested that the opening of chromatin accessibility profiles is correlated with the demethylation of H3K27me3 from E14.5 and during fibroblast lineage commitment. This finding confirms the epigenetic regulation of stem cell differentiation where the PRC2 complex is needed to tri-methylate H3K27 and repress genes required for cellular differentiation (Boyer et al., 2006; Ezhkova et al., 2009; Lee et al., 2006; Thulabandu et al., 2018; Wang et al., 2010). Our model suggests that after Ezh2 establishes an epigenetic profile of early dermal fibroblast progenitors that requires Kdm6b/Jmjd3 activity to initiate chromatin de-repression at differentiation genes.

### Mechanisms regulating adipogenesis and dermal papilla lineage commitment

Lineage tracing and single-cell studies have demonstrated the importance of fibroblast lineages commitment of DFPs in skin development and homeostasis (Driskell and Watt, 2015; Griffin et al., 2020; Plikus et al., 2021). Genetic ablation of Jmdj3/Kmd6b in dermal fibroblasts revealed an inhibition of DFPs to differentiate into terminal fates such as functional Dermal Papilla and Adipocytes. Interestingly, the fibroblast states of papillary and reticular lineages appeared to be unaffected. We found that even though hair follicle morphogenesis was initiated and that Dermal Papilla were observed, Kdm6b-cKO hair follicles appeared stunt and underdeveloped (Figure 4C). Our single-cell RNA-seq experiment comparing WT and Kdm6b-cKO skin revealed a major decrease of Pre-adipocytes in Kdm6b-cKObut not Dermal Papilla (Figure 4I). It is possible that the existence of other epigenetic regulators and histone marks such as bivalency with H3K4me3, or the acetylation of H3K27me3, may regulate the formation but not the function of Dermal Papilla. Our scRNA-seq comparing WT and Kdm6b-cKO skin also detected the significant upregulation of Chromatin Modification GO process, indicating a compensation mechanism to remodel and program the chromatin profile bypassing H3K27me3 demethylation (Figure S2D-E). Our H3K27me3 ChIP-seq evaluating E18.5 WT and Kdm6b-cKOfibroblasts validated the ablation of Kdm6b/Jmdj3 which led to the increase in H3K27me3 levels across the genome. Motif analysis of significantly increased H3K27me3 peaks in Kdm6b-cKOfibroblasts revealed an array of transcription factors that might possess dual-functions. These transcription factors such as Ebf1, Zfp423, Tcf7l2 are not distinctly expressed across fibroblast lineages, even though they have been shown to play important roles in adipogenesis and Dermal Papilla function (Festa et al., 2011; Gupta et al., 2010; Lien et al., 2014). After the integration of scATAC-seq data with our ChIPseq data, we discovered that these transcription factors might regulate different sets of downstream genes in different cell types (Figure 6B). Overall, our data suggested that the chromatin accessibility profiles can dictate the potential of fibroblast commitment to the terminal fates, and that Kdm6b/Jmdj3 demethylation regulates this process. The differential activity of Kdm6b/Jmjd3 in diverse fibroblast lineages acts as a mechanism to regulate the dual-function of non-unique transcription factors. Finally, we have also reconstructed a high-resolution molecular blueprint of dermal fibroblast differentiation processes across multiple modalities including transcriptomic, accessible chromatin profiles, and epigenetic markers to define DFPs and distinct fibroblast fates, freely accessible at [our webtool](https://skinregeneration.org/).

### scATAC-seq offered more insights into the regulation of progenitor cell differentiation and adult tissue homeostasis

Recent studies have characterized the single-cell chromatin profiles of fibroblasts in murine skin wound healing. In regular scarring wounds, Foster and colleagues found different populations of fibroblasts with distinct putative roles (Foster et al., 2021). While comparing large WIHN and small scarring wounds, another study defined the distinct chromatin landscapes of regenerative and scarring fibroblasts (Abbasi et al., 2020), although the competent regenerative fibroblasts might arise from the periphery regenerative fibroblasts (Phan et al., 2021). Biernaskie and colleagues have also discovered the inaccessible inflammatory regulome in reindeer fibroblasts which could lead to the regenerative states found in reindeer’s velvet (Sinha et al., 2022). However, the scATAC-seq datasets were limited by the low cell numbers which could lead to an ineffective interpretation of fibroblast heterogeneity. Altogether, these studies indicated that the dynamic cellular state of dermal fibroblasts is defined by accessible chromatin profiles, and that manipulation of distinct regulatory mechanisms to achieve a regenerative fibroblast state might hold the key to promoting scarless wound healing. Alternatively, there may not be an advantage for achieving a regenerative fibroblast state but to utilize and achieve optimal populations of different committed fibroblast populations that arise from their distinct lineages. Our current study provides an in-depth analysis looking at the changes in chromatin architecture happening during embryonic development while establishing the ground state of accessible chromatin, transcriptomic, and epigenetic profiles for DFPs and differentiating fibroblasts.

In the context of aging, dermal fibroblasts have been shown to have lost their cellular identity at transcriptomic and epigenetic levels (López-Otín et al., 2013; Salzer et al., 2018; Yang et al., 2023). Future studies will requireS the identification of the homeostatic chromatin states in different fibroblast populations to activate fibroblasts to resist the effect of chronic aging. Our molecular blueprint will provide the reference points of differentiating fibroblasts in embryogenesis and postnatal dermis to help determine the chromatin profiles of how fibroblast heterogeneity was established.

## MATERIALS AND METHODS

### Mouse models

All animal procedures in this study were in accordance with protocols approved by Washington State University Institutional Animal Care and Use Committee ASAF #6723,#6724,#6726. Mice were derived from C57BL/6 background. The following transgenic mouse lines were used in this study: Twist2-Cre (RRID:MGI:3840442), Kdm6b^fl/fl^ (RRID:IMSR_JAX:029615), ROSA26^mT/mG^ (RRID:IMSR_JAX:037456)

### Single-cell suspension digestion

Tissues were collected from embryonic E14.5, E17.5/E18.5, and P5 mice skins. Cells isolation procedure described previously was followed to obtain single-cell suspension from tissues(Jensen et al., 2010).

### Histological analysis

Tissues were collected from mice back skins. For formalin-fixed paraffin-embedded, tissues were fixed with 4% paraformaldehyde in 1X PBS overnight. Hematoxylin and Eosin staining were done with 5 μm-thick sections. Colored images were taken with Nikon E600 and Nikon DS-Fi3 camera. Horizontal wholemount preparation was done as previously published (Salz and Driskell, 2017). Immunofluorescent images were taken with Leica SP8 confocal microscope at 20X objective. Analysis and quantification of H&E images were given unique ID and scored by a blinded researcher.

### Chamber allografting assay

This procedure was performed as previously described (Jensen et al., 2010). 8 x 10^6^ WT neonatal epidermal cells were combined with 10^6^ dermal cells from different timepoints or from different transgenic lines. Combination of cells were injected into silicone chambers grafted on 6 mm circular dorsal wounds in Foxn1^-/-^ mice. Collection of grafted areas were done 28 days post-surgery.

### Single-cell Gene Expression

Single-cell suspensions from E14.5, E17.5, P5 were processed separately for 10x Genomics Single Cell 3’ Gene Expression (v2) PN-120237, targeting 10,000 cells per library. Libraries were sequenced on Illumina HiSeq 4000 (100bp Paired-End). Fastq files were aligned to mm10 reference genome using CellRanger version 6.0.0.

Single-cell suspensions of E18.5 WT and Kdm6bKO dermal preparations were processed separately for 10x Genomics Single Cell 3’ Gene Expression kit (V3.1) PN-1000128, targeting 10,000 cells per library. Libraries were sequenced on Illumina NovaSeq6000 (100 Paired-End). Fastq files were aligned to mm10 reference genome using CellRanger version 6.0.0. Loom files were generated from CellRanger alignment outputs via velocyto packages for RNA velocity analysis(La Manno et al., 2018). Analysis of scRNA-seq data were done using Scanpy and ScVelo (Wolf et al., 2018, Bergen et al., 2020).

Low quality and doublet cells were filtered by larger than 5000 genes or less than 700 genes, less than 10 percent mitochondrial genes. Data were normalized to 10,000 reads per cells, log transformed, then scaled to maximum standard deviation 10. Integration and batch-correction by Time were done using Scanorama (Hie et al., 2019). Differential gene expression was calculated using Diffxpy (https://github.com/theislab/diffxpy).

Partition-based graph abstraction (PAGA) was used to infer the trajectory of continuous cell transitions while preserving the global topology of data (Wolf et al., 2019). RNA velocity analysis was done using dynamical model to retrieve latent time (Bergen et al., 2020).

All codes used to analyze this data is available on our github [page](https://github.com/DriskellLab).

### Single-cell Assay for Transposase-Accessible Chromatin

Nuclei isolation from single cell suspension were prepared targeting 10,000 nuclei per library. Isolated nuclei were processed using 10X Genomics Single Cell ATAC v2 (CG000496). Libraries were sequenced on Illumina NovaSeq 6000 (100bp Paired-end). Fastq files were aligned to mm10 reference genome using Cellranger-atac version 2.1.0. Cellranger outputs including Peaks called, Fragments, and metadata were used for downstream analysis with Signac in R (Stuart et al., 2021). Basic filtering and pre-processing were done screening for Peak region fragment <100000 and Percent reads in peaks > 40%. Clustering analysis was done using latent semantic indexing (LSI) (Cusanovich et al., 2015), dimension reduction was carried out by UMAP (McInnes et al., 2018). scRNA-seq integration with scATAC-seq data for cell type classification was done by Signac Label Transferred methods using scRNA-seq fibroblasts object and scATAC-seq fibroblast object (Stuart et al., 2021).

To improve the peaks calling sensitivity and accuracy, we performed Peak Call within Signac using MACS2 function (Zhang et al., 2008). Differential accessible peaks between clusters were calculated using FindMarkers() function within Signac. Motif binding sites analysis of specific cell type was done using MotifMatchR (https://greenleaflab.github.io/motifmatchr/).

Codes used to analyze this data is available on our github [page](https://github.com/DriskellLab).

### Chromatin Immunoprecipitation Sequencing (ChIP-Seq)

Dermal preparation of E14.5, E17.5 WT; E18.5 WT and Kdm6b-cKO murine skins were expanded in culture to reach 2 x 10^7^ cells in AmnioMAX complete medium. ChIP assay was done using SimpleChIP Plus Enzymatic Chromatin IP kit (magnetic beads) #9005 from CellSignalling and H3K27me3 antibody (Active Motif RRID:AB_2561020). ChIP samples were prepared with Kapa Hyper prep kit. Libraries were sequenced on Illumina NovaSeq 6000 (100bp Paired-End). Fastq ChIP-seq files were aligned using bowtie2 and converted to sorted bam files and BED files. E18.5 WT and Kdm6bKO samples were subsampled to equivalent read depths as other samples (6 x 10^7^ reads per sample). Peak called was done using MACS2 with --broad option and –broad-cutoff q-value =0.01. Visualization of ChIP-seq data was done using ChIPseeker for CoveragePlot (Wang et al., 2022), Easeq for Tracksplot (Lerdrup and Hansen, 2020). Differential ChIP-seq calculation was done using DiffBind (Ross-Innes et al., 2012; Stark and Brown, n.d.).

Motif Analysis was done using HOMER findMotifs function (Heinz et al., 2010).

### Flow cytometry

E18.5 single-cell suspension from WT and Kdm6bcKO mice were stained with primary antibody and secondary antibody for 1 hour each with 3X PBS washes in between and after. Flow sorting of cells were done on Sony SH800 Flow Sorter. Flow analysis were done on ThermoFisher Attune Flow Cytometer.

### Statistical analysis

Histology data comparing two groups were tested for normality using Shapiro-Wilk, equal distribution using Levene, and t-test for independent samples if assumptions are met, otherwise we used Wilcoxon Rank Sum test. Single-cell RNA and ATAC data statistical tests were done using non-parametric Wilcoxon Rank Sum test.

## ACKNOWLEDGEMENTS

This work was supported by NIH National Institute of Arthritis and Musculoskeletal and Skin Diseases (NIAMS) R01 AR078743. We would also like to acknowledge the Franceschi Microscopy & Imaging Center at Washington State University; Center for Reproductive Biology (Melissa Oatley); Gene Editing Reagent Core at Washington State University (Lisette Maddison, Deqiang Miao); Washington State University Functional Genomics Initiative; This publication includes data generated at the UC San Diego IGM Genomics Center utilizing an Illumina NovaSeq 6000 that was purchased with funding from a National Institutes of Health SIG grant (#S10 OD026929); Genomics & Cell Characterization Core Facility (GC3F) at University of Oregon; Laboratory for Biotechnology and Bioanalysis at Washington State University (Derek Pouchnik, Mark Wildung, and Weiwei Du), QMP, JM was supported by a National Institute of General Medicine Sciences training grant (T32GM008336), QMP was supported by Poncin Fellowship.

## EXPANDED FIGURE LEGENDS

**Figure S1.**
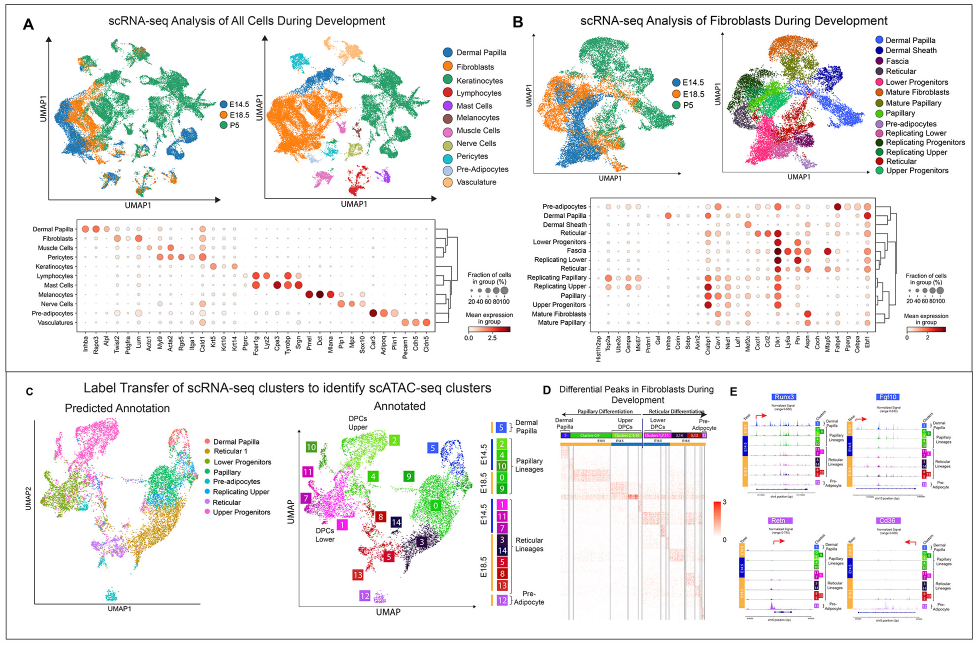
scRNA-seq and scATAC-seq analysis. (A) scRNA-seq analysis of all cells from E14.5, E18.5, and P5 skin. (B) scRNA-seq analysis of fibroblast subset re-clustered from E14.5, E18.5, and P5 skin. (C) Label transfer experiment of scRNA-seq cluster to identify scATAC-seq cluster in E14.5 and E18.5 skin. (D) Differential peak analysis of scATAC-seq clusters arranged in pseudotime. (E) Trackplot of fibroblast clusters arranged in pseudotime for Runx3, Fgf10, Retn, and CD36.

**Figure S2.**
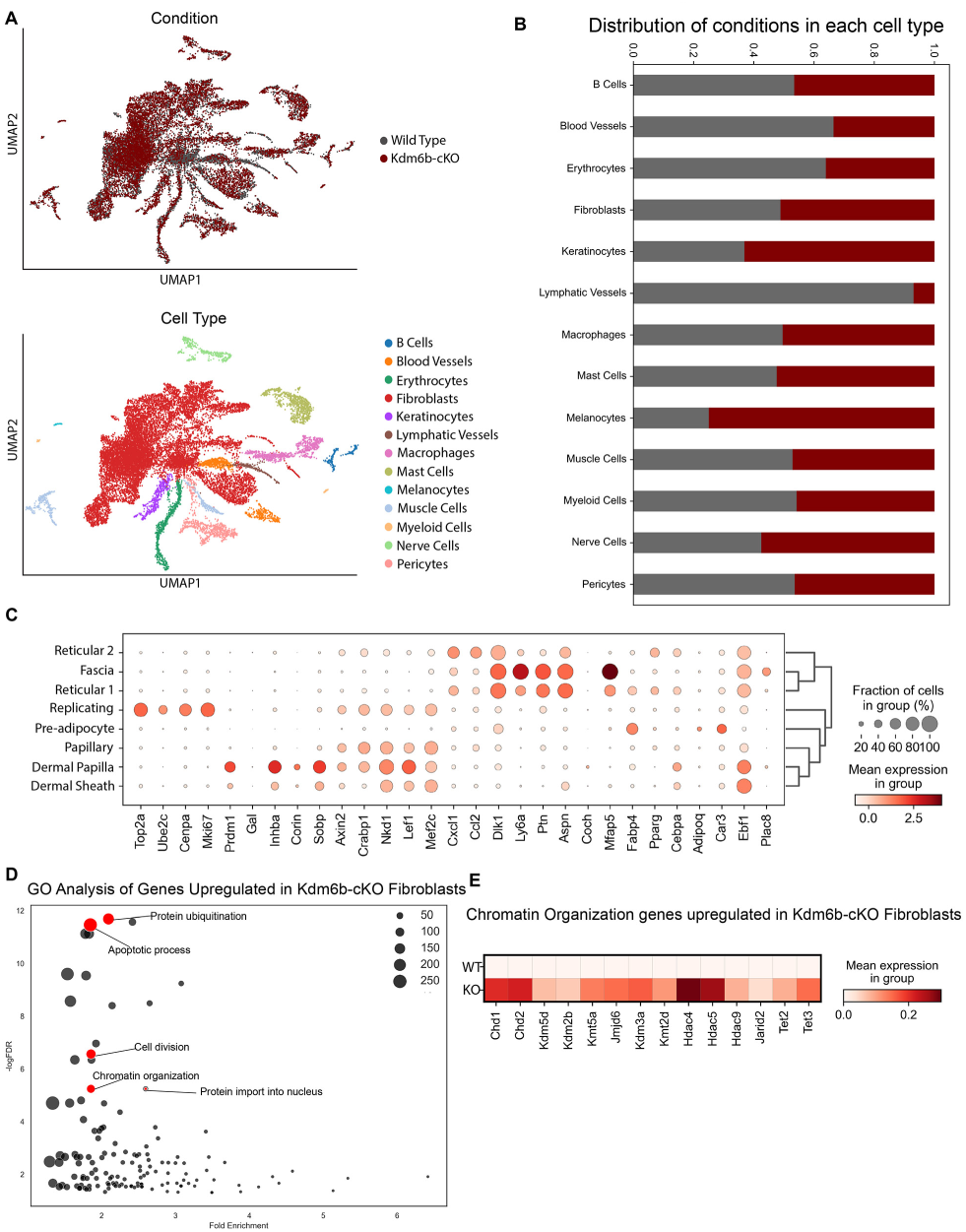
scRNA-seq analysis of all cells in WT and Dermo1Cre-Kdm6b^fl/fl^ skin. (A) UMAP projections of WT and Dermo1Cre-Kdm6b^fl/fl^ skin. (B) Quantification of the number of cells within each cluster type based on genotype represented as a percentage in the cluster. (C) Dotplot of gene utilized to identify cell clusters. (D) GO analysis of differentially regulated genes comparing WT and Dermo1Cre-Kdm6b^fl/fl^ skin. (E) Gene expression of Chromatin Organization genes from GO Analysis.

**Figure S3.**
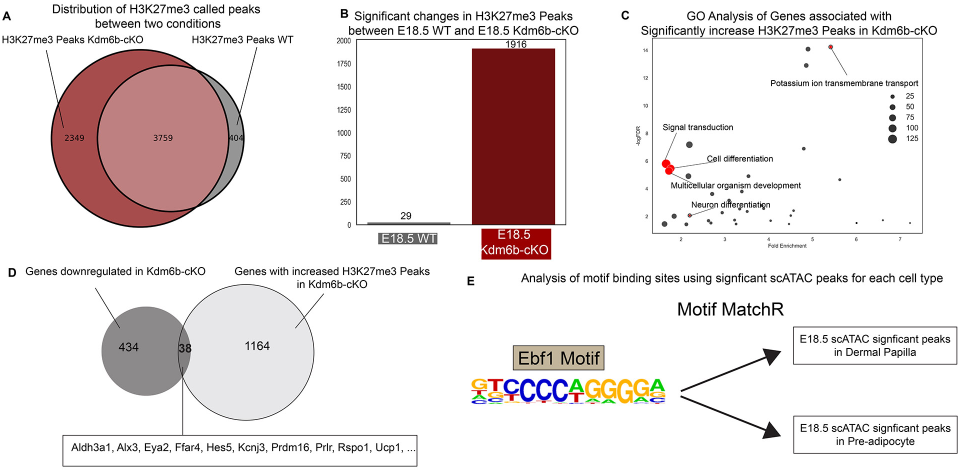
Integrative analysis of H3K27-me3 peaks between WT and Dermo1Cre-Kdm6b^fl/fl^ fibroblasts with integrated motif analysis in scATAC data and scRNA-seq data. (A) Venn diagram comparing H3K27me3 peaks between WT and Dermo1Cre-Kdm6b^fl/fl^ fibroblasts. (B) Quantitation of significantly called peaks from WT and Dermo1Cre-Kdm6b^fl/fl^ fibroblasts. (C) GO Analysis of peaks significantly increased in Dermo1Cre-Kdm6b^fl/fl^ fibroblasts. (D) Venn diagram comparing downregulated genes in scRNA-seq analysis and peaks in ChIPseq analysis. (E) Motif analysis of scATAC peaks for GO analysis.

